# Altered cortical brain structure and increased risk for disease seen decades after perinatal exposure to maternal smoking: A study of 9,000 adults in the UK Biobank

**DOI:** 10.1101/471839

**Authors:** Lauren E. Salminen, Rand R. Wilcox, Alyssa H. Zhu, Brandalyn C. Riedel, Christopher R. K. Ching, Faisal Rashid, Sophia I. Thomopoulos, Arvin Saremi, Marc B. Harrison, Anjanibhargavi Ragothaman, Victoria Knight, Christina P. Boyle, Sarah E. Medland, Paul M. Thompson, Neda Jahanshad

## Abstract

Secondhand smoke exposure is a major public health risk that is especially harmful to the developing brain, but it is unclear if early exposure affects brain structure during middle age and older adulthood. Here we analyzed brain MRI data from the UK Biobank in a population-based sample of individuals (ages 44-80) who were exposed (n=2,510) or unexposed (n=6,079) to smoking around birth. We used robust statistical models, including quantile regressions, to test the effect of perinatal smoke exposure (PSE) on cortical surface area (SA), thickness, and subcortical volume. We hypothesized that PSE would be associated with cortical disruption in primary sensory areas compared to unexposed (PSE^-^) adults. After adjusting for multiple comparisons, SA was significantly lower in the pericalcarine (PCAL), inferior parietal (IPL), and regions of the temporal and frontal cortex of PSE^+^ adults; these abnormalities were associated with increased risk for several diseases, including circulatory and endocrine conditions. Sensitivity analyses conducted in a hold-out group of healthy participants (exposed, n=109, unexposed, n=315) replicated the effect of PSE on SA in the PCAL and IPL. Collectively our results show a negative, long term effect of PSE on sensory cortices that may increase risk for disease later in life.

## Introduction

Tobacco use is a major public health concern worldwide. According to the World Health Organization (WHO), there are currently 1.1 billion daily smokers globally, exposing more than one third of the adult world population to secondhand smoke. Global smoking trends show a recent net decline in smoking prevalence, but this is primarily driven by decreased smoking prevalence in high income countries. Further, the total number of smokers worldwide has not fallen overall due to population growth, particularly in low- and middle-income countries (LMICs) that account for 80% of the world’s daily smokers (WHO 2018). Tobacco smoke is currently responsible for 6 million deaths annually, including 600,000 deaths from secondhand smoke. If current trends persist, around half of current smokers will die prematurely from tobacco use by 2050, with a disproportionate hit to LMICs (WHO 2018).

Health risks associated with cigarette smoke can be traced to more than 7,000 chemical compounds, including over 4,500 toxins and approximately 70 carcinogens (Choi et al. 2017) Approximately 15% of cigarette smoke is ingested by the smoker, while the remainder is released into the atmosphere. Cigarette smoke is more harmful to children, infants, and the developing fetus than to adults due to their faster respiratory rates that result in greater smoke consumption. Their underdeveloped lungs and immune systems further increase the risk for acute and chronic illnesses (CDC, 2018). Accordingly, exposure to maternal smoking and environmental tobacco smoke increases risk for spontaneous preterm abortion, poor intrauterine lung function, low birthweight in neonates (Hofhuis et al. 2003), Sudden Infant Death Syndrome (SIDS), middle ear disease, asthma and respiratory disease, vision problems, neurodevelopmental conditions in children, cardiovascular disease, and psychiatric illness (Polanska et al. 2008; Blood-Siegfried and Rende 2010; CDC 2017). There is also increasing evidence that perinatal smoke exposure (PSE) influences structural brain development (Toro et al. 2008; Derauf et al. 2012; Liu et al. 2013; El Marroun et al. 2014), which may impair physical and mental health later in life.

Nicotine is the most widely studied constituent of tobacco smoke due to its addictive properties and ability to enter the brain within seconds of consumption (Berridge et al. 2010). In the brain, nicotine alters the activity of nicotinic acetylcholine receptors (nAChRs), which play an important role in neuronal maturation when bound by the neurotransmitter, acetylcholine (ACh). The pharmacological action of nAChRs is determined by homomeric and heteromeric complexes of α and β protein subunits that are widely distributed throughout the brain (Gotti et al. 2006). Localized expression of nAChR subunit configurations facilitates distinct functions during early brain development, including (but not limited to) neuronal pathfinding, the formation of essential neural circuits and sensory structures (Bansal et al. 2000) and regulation of brain stem networks (O’Leary et al. 2008). In the early postnatal period, the functional patterns of nAChRs are critical for regulating synaptic pruning in the neocortex (Orr-Urtreger et al. 2000), thalamocortical synaptic maturation, and sensory signaling (Metherate and Hsieh 2003; Aztiria et al. 2004). Accordingly, dysregulated nAChR activity in animal studies and wet lab work has been linked to attenuated neuronal differentiation, apoptosis, desensitized synaptic circuits, and lower cell density in the somatosensory cortex (Slotkin 2004; Chatterton et al. 2017). Other work (Swan and Lessov-Schlaggar 2007; Coggins et al. 2013; Singh et al. 2017) suggests there is a multi-mechanistic role of nicotine and other harmful cigarette compounds on brain structure, but human research in this area is limited relative to articles on the bioactive effects of nicotine.

Most tobacco-related morbidities occur decades after initial tobacco use and secondhand smoke exposure (Holford 2014). Thus, today’s current population of middle-aged individuals (ages 40-65) is the highest risk group for tobacco-related illness and early death. This mortality risk includes non-smokers exposed to secondhand smoke, with disproportionately higher risk among those exposed at younger ages (Diver et al. 2018). PSE has been identified as a long-term risk for cancer, cardiovascular disease, and other systemic morbidities (CDC 2017), but we lack understanding of the long-term effects of PSE on brain structure in middle and old age, and how this altered brain structure may further relate to risk for health conditions. Given the known biological repercussions of tobacco smoke and delayed onset of smoke-attributable disease, identifying the long-term effects of PSE on brain structure may be used to inform tobacco control policies and public health warnings for smoking prevention and cessation (WHO 2018).

The UK Biobank (UKB) is currently the largest prospective study of aging, collecting detailed health, physical, and lifestyle information for 500,000 middle aged and older adults living in the United Kingdom (UK). Here we analyzed MRI data from the UKB to examine cortical and subcortical brain structure in a subset of PSE^+^ and PSE^-^ (unexposed) individuals who completed neuroimaging. Based on earlier work (Toro et al. 2008; Derauf et al. 2012; Liu et al. 2013; El Marroun et al. 2014), we hypothesized that cortical measurements would be preferentially associated with PSE relative to subcortical gray matter indices, with the strongest effects in the primary sensory cortices - regions that mature rapidly during the neonatal period.

## Methods

### Participants

Data were analyzed from a population-based sample of individuals from the UKB (Application ID#11559; July 2017 release) who completed neuroimaging (N=8,589; males, n=4,050, females, n=4,539; *M*_age_= 62.5, *SD*=7.5), and a separate group of participants without any self-reported or medically documented health conditions (herein referred to as “elite healthy” (N=424; males, n=218, females, n=206; *M*_age_= 59.7, *SD*=7.2). The impetus to examine elite healthy individuals separate from the full sample was to better isolate effects of PSE by eliminating individuals with conditions that might directly or indirectly impact brain structure. All participants were between the ages of 44 and 80 years. Exclusion criteria for the whole sample were limited to safety factors that precluded neuroimaging (e.g., metal implants, recent surgery, etc.). Individuals in the elite healthy sample were excluded for any documented health condition identified through self-report or hospital records indexing medical diagnoses via the 10^th^ revision of the International Statistical Classification of Diseases and Related Health Problems (ICD-10), and contraindications for MRI. All participants provided informed consent prior to study participation. Data analyzed in this manuscript are publicly available via approved research applications submitted to UK Biobank (http://www.ukbiobank.ac.uk/register-apply/).

### Neuroimaging acquisition

Participants completed a 31-minute neuroimaging protocol at the UKB imaging center in Cheadle, Manchester, UK. All scans were conducted on a single Siemens Skyra 3 tesla scanner, running software VD13A SP4 with a 32-channel RF receive head coil. Structural T1-weighted brain scans were acquired using the following parameters: 3D MPRAGE, sagittal orientation, inplane acceleration factor=2, resolution=1.0 x 1.0 x 1.0 mm^3^ voxels, TI/TR=800/2000 ms. Scans were pre-scan normalized using an on-scanner bias-field correction filter. The scanning protocol is further detailed in Miller et al. (2016) and Alfaro-Almagro et al. (2018).

### Image processing

At the time of the July 2017 data release and analysis, cortical surface area (SA) and thickness (CT) were not available from the UKB data showcase. We therefore, independently extracted CT and SA values using FreeSurfer version 5.3 (Fischl 2012) and the cortical extraction and quality control (QC) protocols we have developed for the ENIGMA consortium (http://enigma.usc.edu/protocols/imaging-protocols/). The extraction protocol yields CT and SA values for 68 cortical regions of interest (ROIs) across the left and right hemispheres according to the Desikan-Killiany atlas (Desikan et al. 2006). FreeSurfer segmentations underwent a rigorous QC procedure, detailed on the ENIGMA website. Participants with scans that failed the visual QC entirely, due to either anatomical abnormalities or scan artifacts, were excluded. Out of 10,573 scans, 171 failed the visual QC entirely and were unusable. Of the remaining scans, 3,412 exhibited minor segmentation errors in which values for discrete regions were removed prior to statistical analysis. As we did not hypothesize any lateralized effects of PSE, measures from bilateral ROIs were averaged across hemispheres for data reduction.

We used the imaging-derived phenotypes released by UKB to examine subcortical volumes (VOL) for the thalamus, pallidum, putamen, caudate, amygdala, hippocampus, and nucleus accumbens, which were computed using *FMRIB’s Integrated Registration and Segmentation Tool* (FIRST; Patenaude et al. 2011) and averaged across both hemispheres. *Operational definitions of predictor variables*

#### Perinatal smoke exposure (PSE)

PSE was assessed from a touchscreen questionnaire on early life factors. Participants were asked, “Did your mother smoke regularly around the time when you were born?” (data field 1787, http://biobank.ctsu.ox.ac.uk/crystal/field.cgi?id=1787). Responses were recorded as “yes” (n=2,663), “no” (n=6,502), “do not know” (n=552), or “prefer not to answer” (n=1). We analyzed “yes” (exposed) and “no” (unexposed) responses.

#### Sociodemographic factors

Proxy measures for sociodemographic factors were examined to identify potential confounders of brain structure that are salient in PSE^+^ populations. The following variables were examined in addition to age and sex:

<u>Educational attainment</u> was assessed using a sociodemographic questionnaire in which participants reported their completed qualification levels according to the UK education system (data field 6138). Possible responses included^1^ “College or University Degree”, “A levels/AS levels or equivalent”, “O levels/GCSEs or equivalent”, “CSEs or equivalent”, “NVQ or HND or HNC or equivalent”, “Other professional qualification”, “None of the above”, or “Prefer not to answer”. To generalize results beyond the UK education system, we converted each response to the corresponding “number of years” using ISCED harmonization guidelines (Goujon et al. 2016). Closer examination of these variables revealed a relatively small sample space for years of education, which functioned as a discrete variable in our dataset. To improve the stability of our statistical models, we dichotomized education into ‘college’ versus ‘high school’ education groups based on the maximum number of completed education levels (≥17 yrs.=college according to the UK education system).

<u>Socioeconomic status (SES)</u> was measured using the Townsend Index (Townsend 1988) (data field 189) – a measure of societal deprivation – based on the postal code of the participant. Townsend scores are widely used as a proxy measure of SES (Smith et al. 2001); prior work in this sample shows high genetic correlations between Townsend scores, psychiatric conditions, and diseases of the central nervous system, as well as specific biomarkers of physical health (Hill et al. 2016). Townsend scores were multiplied by -1 so that higher scores indicated higher SES.

<u>Population structure</u> was assessed using four principal components from the multidimensional scaling protocol of the UKB genetic ancestry analysis.

#### Adult Smoking

We examined potential intervening effects of adulthood smoking on brain measures using the variable “Smoking status” (data field 20116); responses included never smoked” (n=5,670), “previous smoker” (n=3,116), “current smoker” (n=362), or “prefer not to answer” (n=17). To reduce the number of comparisons, we collapsed previous and current smokers into one group and contrasted them against “never smokers” in the main analyses.

#### Waist-to-hip ratio (WHR)

We analyzed WHR (data fields 48-49) to index potential differences in physical/cardiovascular health between PSE groups. WHR is a common index of central fat distribution that has been linked to PSE (Santos et al. 2016). Prior work shows a strong positive correlation between WHR and visceral fat in healthy adults (Gadekar et al. 2018). WHR is also a marker of atherosclerotic burden in overweight individuals and postmenopausal women (Lee et al. 2015; Scicali et al. 2018), and a risk factor for major cardiovascular events and death in women with heart disease (Medina-Inojosa et al. 2018; Streng et al. 2018), and mortality in men (Mousavi et al. 2015). Waist and hip measurements were obtained manually at the time of the imaging scan.

<u>Birth weight</u>. Birth weight was reported retrospectively in 5,915 participants of the population-based sample and 292 participants in the elite healthy sample. Maternal smoking during pregnancy is consistently linked to low birthweight in neonates if smoking behaviors continue beyond the first trimester (Ding et al. 2017). Thus, we analyzed whether observed effects between PSE and brain structure would persist after covarying for differences in birth weight – an indicator of prenatal health (Gortmaker 1979; Kogan et al. 1994; Alexander and Korenbrot 1995). Birth weight was reported in kg units.

### Analytic approach

Chi-squared analyses and independent samples *t*-tests were run to describe potential differences in sociodemographics, adult smoking behavior, and WHR between PSE^+^ (perinatally exposed) and PSE^-^ (unexposed) groups. These variables served as covariates in the main analyses and did not indicate problematic multicollinearity indexed with the variance inflation factor. Continuous covariates and dependent variables were *z*-transformed prior to analysis to standardize results and facilitate interpretation. We also performed chi-squared tests between PSE and binary ICD-10 codes to better characterize the population-based sample. As the cell sizes for many conditions were limited, we aggregated individual disease codes into ICD-10 chapter codes using the ICD classification system (https://www.icd10data.com/ICD10CM/Codes).

For the primary analyses, we computed three series of ordinary least squares (OLS) linear regression models to determine whether brain structure differed between PSE^+/-^ groups using imaging metrics of (1) cortical thickness (CT), (2) cortical surface area (SA), and (3) subcortical volumes (VOL). Age and sex were included as covariates in all models, and intracranial volume (ICV) was included as a covariate in SA and VOL regressions. False discovery rate (FDR) corrections were applied to correct for multiple comparisons across the 75 brain metrics (34 CT metrics, 34 SA metrics, 7 VOL metrics) using the Benjamini-Hochberg method (Benjamini and Hochberg 1995). Regression models were then re-analyzed using the same approach as the main analysis after including birth weight as a covariate predictor. We did not covary for birth weight in the main analysis to avoid regressing out the effects of PSE and/or conflating them with other prenatal factors (stress, nutrition, etc.).

Sensitivity analyses were completed in the elite healthy sample (PSE^+^, n=109; PSE^-^, n=315) for brain regions that differed significantly by PSE using the same statistical analyses in the population-based sample. These participants were held out of main analyses for replication.

### Assessing heteroscedasticity

Neuroimaging variables were checked for normality and homoscedasticity at the model level using Kolmogorov-Smirnov (Massey 1952) and Koenker-Bassett tests (Koenker and Bassett 1982). Outliers and leverage points among the covariates were checked using robust projection methods described by Wilcox (Wilcox and Keselman 2012) via the ‘WRS2’ package in R (Mair and Wilcox 2017). Examination of the data indicated there was heteroscedasticity. That is, the variation in the neuroimaging variables depended on the values of the independent variables. Additionally, the data were not normal. We used two methods to combat these issues. First, we applied an HC4 standard error estimator to the OLS models using the `‘olshc4’` function in R. The HC4 method (Cribari-Neto 2004) deals with heteroscedasticity data but is highly conservative, with a null rejection rate at ~1% (Cribari-Neto and Maria da Gloria 2014). Moreover, OLS can be negatively impacted by outliers. Thus, we computed quantile regressions in addition to OLS models to examine neuroimaging differences between PSE groups. To do this, we used the ‘`regci`’ function in R (Wilcox and Keselman 2012) with a quantile regression estimator. To provide a more detailed understanding of the association, we tested the hypothesis that predictor slopes were *not* equal to zero using the (conditional) quantiles = 0.1, 0.25, 0.5, 0.75, 0.9 associated with the dependent variable. We also applied a separate FDR correction threshold for the 75 metrics across each quantile (375 comparisons).

When comparing groups, effect sizes are reported using robust analogs of Cohen’s *d* using the R function `‘akp.effect’` (Algina et al. 2005), which is based on 20% trimmed means and Winsorized standard deviations. Effect sizes were computed for each Winsorized standard deviation when groups had unequal variances. The magnitude of effect size can be interpreted using Cohen’s guidelines: 0.2 (small), 0.5 (medium), and 0.8 (large). We also report *explanatory power* – an estimate of effect size that is robust to unequal variances between groups (Wilcox and Tian 2011).

To add perspective, effect sizes were computed for brain metrics that differed by PSE using the quantile shift function in R (Wilcox, in press). Briefly, when groups do not differ, the median of the typical difference, *x* - *y*, is zero. A quantile shift effect size (Q) reflects a shift of the median to a higher or lower quantile associated with the distribution of *x-y*; it deals with nonnormality and does not assume homoscedasticity. Under normality and homoscedasticity, Cohen’s *d* = 0, 0.2, 0.5, and 0.8 corresponds approximately to a Q effect of 0.5, 0.55, 0.65, and 0.7, respectively (Wilcox, in press).

### Brain structural contributions to PSE-related disease risk

To determine the clinical relevance of the significant models observed in our study, we completed *post hoc* logistic regressions to determine relative risk of developing specific classes of ICD-10 medical conditions (e.g., psychiatric). We used a Partial F-test to determine whether region-specific brain structure significantly contributed to explaining the population variance. “Partial models” were computed to determine relative risk for each ICD-10 condition class using each independent variable from our main analysis. “Full models” were then computed by adding neuroimaging variables that differed significantly by PSE in the main analysis to each predictor set. These models were first conducted in the whole sample to determine the relative risk of PSE, and then conducted within groups. These analyses were completed in a reduced sample that removed all listwise missingness (PSE^+^ =1,945; PSE^-^ = 4,784). Significance for these analyses was determined at a threshold of *p*<0.05, as these analyses were *post-hoc* and included regions and disorders already determined to be significant after multiple comparisons correction; further correcting for multiple comparisons may have a greater public health risk of a type 2 error. We did not include birth weight as a predictor in these analyses given the lower number of participants who provided this information (N=5,915, further reduced when accounting for each disease), and its inherent association to PSE. However, we did use ICV as a predictor in these models, as smaller head size is a reported outcome of prenatal smoking and early environmental tobacco smoke exposure (Bouthoorn et al. 2012). We did not examine ICD-10 classes of “Symptoms not otherwise specified” (ICD codes R00-R99), “Consequences of external causes” (S00-T88), “External causes of morbidity” (V00-Y99), or “Factors influencing health status” (Z00-Z99) given the heterogeneous nature of these ICD categories.

All analyses were conducted using two-tailed tests in R, version 3.4.

## Results

### Cohort demographics and comparisons of descriptive variables

Sociodemographic characteristics of the sample are provided in **Table 1**. The sample size varied per analysis due to differences in data availability across imaging metrics, ROIs that failed the cortical QC, and presence of outliers. Participants in the PSE^+^ group were significantly younger (*t*(8587)=2.7, *p*=0.006), had a higher WHR (*t*(8578)=-3.5, *p*<0.001), lower education (*t*(8582)=5.11, *p*<0.001), and lower SES indicated by their Townsend scores (*t*(8583)=3.6, *p*<0.001). Chi-squared analysis showed that a greater proportion of the PSE^-^ (unexposed) group was male (*χ^2^*(1)=584.1, *p*<0.001) and a past or current smoker (*χ^2^*(1)=156.7, *p*<0.001) when compared to the PSE^+^ group. Chi-squared analysis also showed that a lower proportion of the PSE^+^ group had completed any years of college when compared to the PSE^-^ group (*χ^2^*(1)=13.9, *p*<0.001). Analysis of ICD-10 conditions showed that a greater proportion of the PSE^+^ group was diagnosed with the following classes of disease: psychiatric (*χ^2^*(1)=11.6, *p*<0.001), central nervous system (*χ^2^*(1)=5.77, *p*=0.016), circulatory (*χ^2^*(1)=6.79, *p=0.009),* respiratory (*χ^2^*(1)=5.69, *p*=0.017), digestive (*χ^2^*(1)=8.97, *p=0.003*), musculoskeletal (*χ^2^*(1)=4.15, *p*=0.042), and genitourinary conditions (*χ^2^*(1)=7.42, *p*=0.006).

**Table 1.** Descriptive Characteristics of the Samples

| Descriptive Variables | Population-based Sample<br>N=8,589 |  | Elite Healthy Replication Sample<br>N=424 |  |
| --- | --- | --- | --- | --- |
|  | PSE<sup>-</sup> | PSE<sup>+</sup> | PSE<sup>-</sup> | PSE<sup>+</sup> |
| Age, M(SD) | 62.7(7.6)** | 62.2(7.1)** | 59.6(7.4) | 59.9(6.4) |
| Waist/hip ratio, M(SD) | 0.86(0.08)*** | 0.87(0.08)*** | 0.85(0.8) | 0.84(0.8) |
| Townsend Index, M(SD) | 2.0(2.6)*** | 1.8(2.7)*** | 2.1(2.5) | 2.3(2.5) |
| Sex, % Male | 47.3% | 46.7% | 47.3% | 52.3% |
| Education, % College | 68.3%*** | 64.1%*** | 76%*** | 64%*** |
| Adult Smoker, % Yes | 37.8%*** | 35.1%*** | 29.5% | 30.3% |
| **ICD-10 Conditions by ICD Chapter <sup>a</sup>, % Yes:** |  |  |  |  |
| Infections/parasitic disease (A00-B99) | 4.7% | 5.4% | -- | -- |
| Neoplasms (C00-D49) | 17.9% | 18.4% | -- | -- |
| Blood (D50-D89) | 3.8% | 4.2% | -- | -- |
| Endocrine (E00-E89) | 12.0% | 12.7% | -- | -- |
| Psychiatric (F01-F99) | 3.8%** | 5.4%** | -- | -- |
| CNS (G00-G99) | 5.9%* | 7.3%* | -- | -- |
| Eye (H00-H59) | 7.9% | 7.8% | -- | -- |
| Ear (H60-H95) | 1.5% | 2.0% | -- | -- |
| Circulatory (I00-I99) | 23.8%** | 26.4%** | -- | -- |
| Respiratory (J00-J99) | 10.6%* | 12.3%* | -- | -- |
| Digestive (K00-K95) | 32.6%** | 36.0%** | -- | -- |
| Skin (L00-L99) | 9.2% | 8.4% | -- | -- |
| Musculoskeletal (M00-M99) | 18.7%* | 20.6%* | -- | -- |
| Genitourinary (N00-N99) | 22.8%** | 25.6%** | -- | -- |
| Pregnancy-related (O00-O9A) | 4.3% | 4.1% | -- | -- |
| Congenital (Q00-Q99) | 0.8% | 1.1% | -- | -- |

Note: Demographic numbers vary slightly per analysis. <sup>a</sup> Not reported are conditions originating in the perinatal period, as there was only one subject in our sample with an ICD-10 diagnosis in this category. We also did not include ICD-10 codes after Q99, as these ICD categories are heterogeneous and challenging to interpret in relation to brain metrics. \*\*\* $p<0.001$ , \*\* $p<0.01$ , \* $p<0.05$

**Figure 1.**
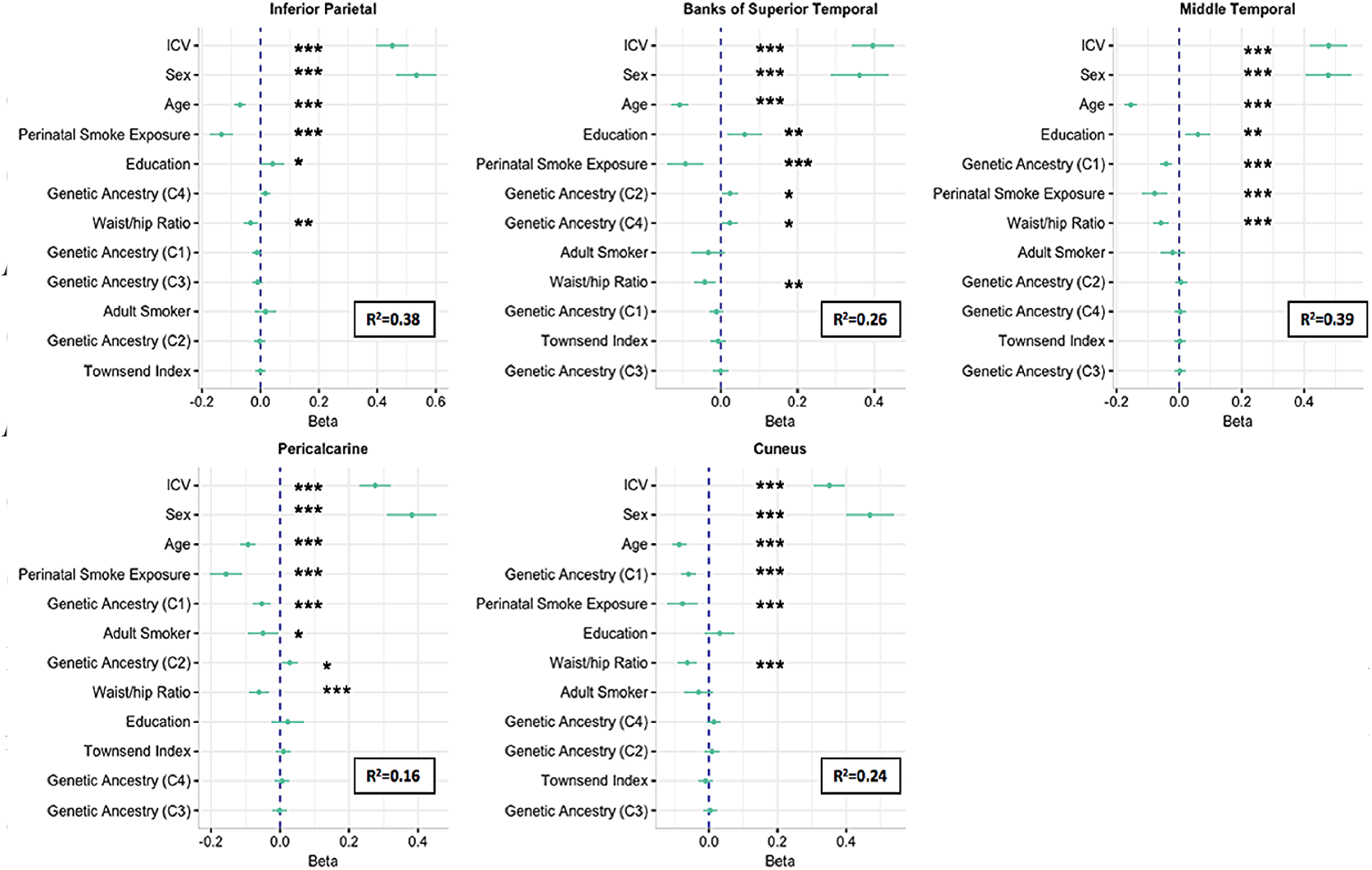
Beta weights for cortical surface area regions that differed by PSE in the population-based sample. Predictor variables are plotted on the y axis in order of importance using the lasso method. Individual principle components of genetic ancestry are indicated in parentheses. ***p<0.001, **p<0.01, *p<0.05.

PSE groups within the elite healthy replication sample did not differ significantly on age (*t*(422)=-0.26, *p*=0.794), WHR (*t*(422)=0.78, *p*=0.433) or Townsend scores (*t*(421)=-0.44, *p*=0.657). Chi-squared tests showed that a smaller proportion of the PSE^+^ group was male (*χ^2^*(1)=58.1, *p*<0.001) and had a college-level education (*χ^2^*(1)=187.9, *p*<0.001). There were no significant group differences in sex or adult smoking (*χ^2^*(1)=1.7, *p*=0.229).

### Brain structural differences by PSE

Quantile regressions revealed similar findings in some regions, but different effects in others. Specifically, individuals in the PSE^+^ group showed significantly lower SA in the PCAL and IPL at quantiles 0.1-0.75, the bSTS at quantiles 0.1-0.5, the MTG at quantiles 0.1-0.25 and 0.75-0.9, and the ITG at quantile 0.25 (**Figure 2a)**. Trend effects (*p*<0.01) were observed for lower SA in the CUN and RMFG at quantile 0.75, and lower VOL in the caudate, pallidum, and putamen at quantile 0.75. The pallidum also trended towards significance at the 0.9 quantile (**Supplementary Table 1**). SA and VOL in these regions were negatively associated with age, and positively associated with sex and ICV at all quantiles of the regressions. After age, sex, and ICV, the two most common predictors of SA were WHR and the first multidimensional scaling component of genetic ancestry (C1). WHR was also negatively associated with VOL in the caudate, pallidum, and putamen, but associations varied across quantiles. Finally, college education was positively associated with VOL in the caudate, pallidum, and putamen, but associations were only observed at lower quantiles (q=0.1-0.5). Quantile shift effect sizes were small in all regions that differed significantly by PSE or trended towards significance (**Table 3a**).

**Figure 2.**
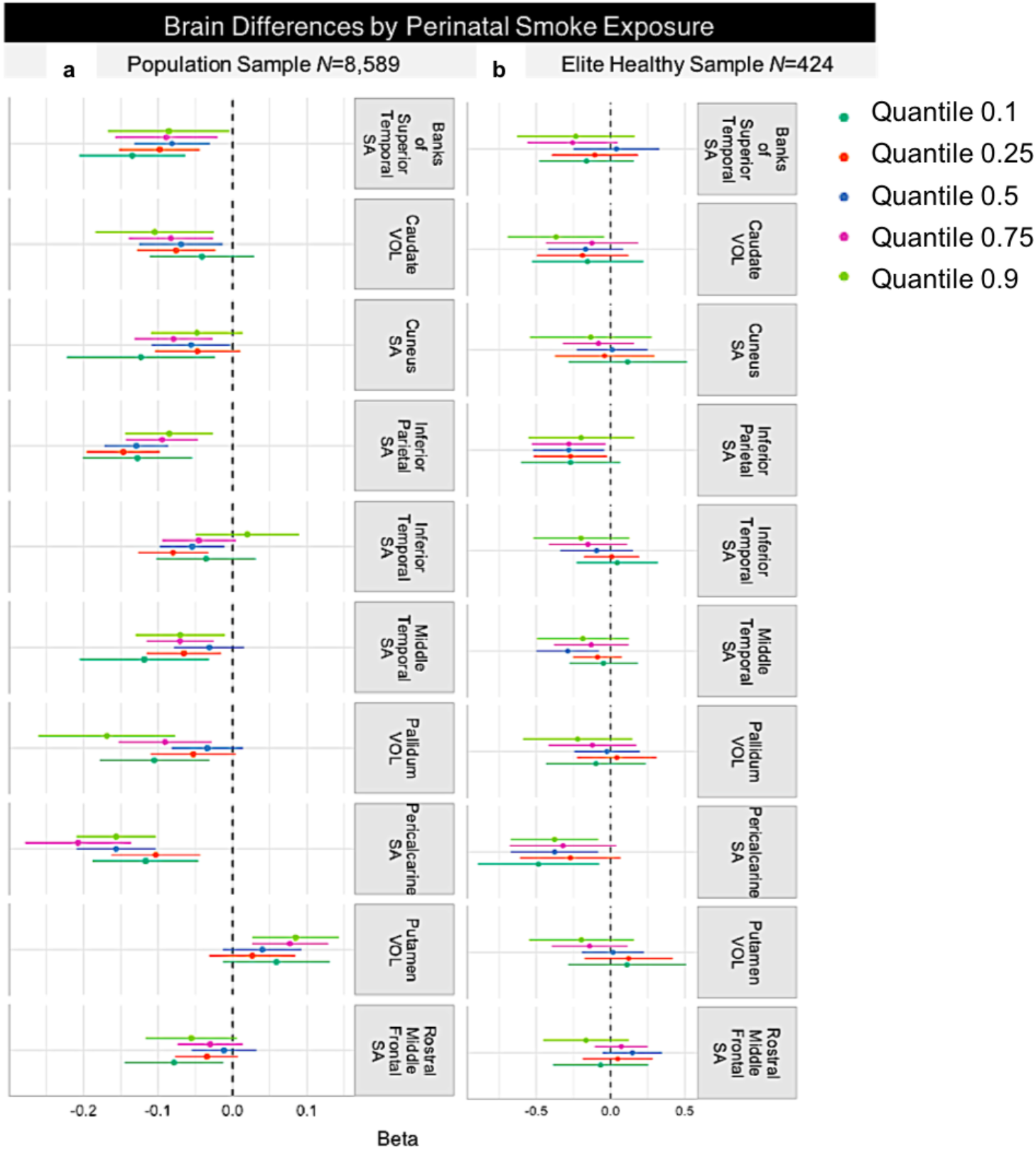
Beta weights of the neuroimaging metrics that differed significantly by PSE. (**a**) Dot-whisker plots on the left depict observed brain differences in the population-based sample. (**b**) Plots on the right reflect the replication results in the elite healthy sample. Only trend level effects were observed in the elite healthy replication sample. SA= surface area, VOL=volume

After covarying for birth weight, the PSE^+^ group demonstrated significantly lower SA in the PCAL and IPL in all quantiles. Effects were strongest for the 0.25 and 0.5 quantiles for the IPL (β’s= -0.1 and -0.12, respectively), and the 0.25-0.75 quantiles for the PCAL. Birth weight was positively associated with SA in the IPL for all quantiles (**Supplementary Table 1**). We did not detect significant associations between birth weight and SA in the PCAL at any point along the distribution (**Supplementary Table 2**).

### Replication in the elite healthy sample

Replication analyses using the OLS approach revealed lower SA in the PCAL (n=354; β=-0.36, 95% CIs[-0.53, -0.07], *p*=0.009), IPL (n=362; β=-0.32, 95% CIs[-0.49, -0.14], *p*<0.001), and MTG (n=341; β=-0.19, 95% CIs[-0.36, -0.02], *p*=0.028) of the PSE^+^ group. We did not replicate the effects of PSE in the bSTS or CUN that were observed using the OLS approach (**Figure 3**).

**Figure 3.**
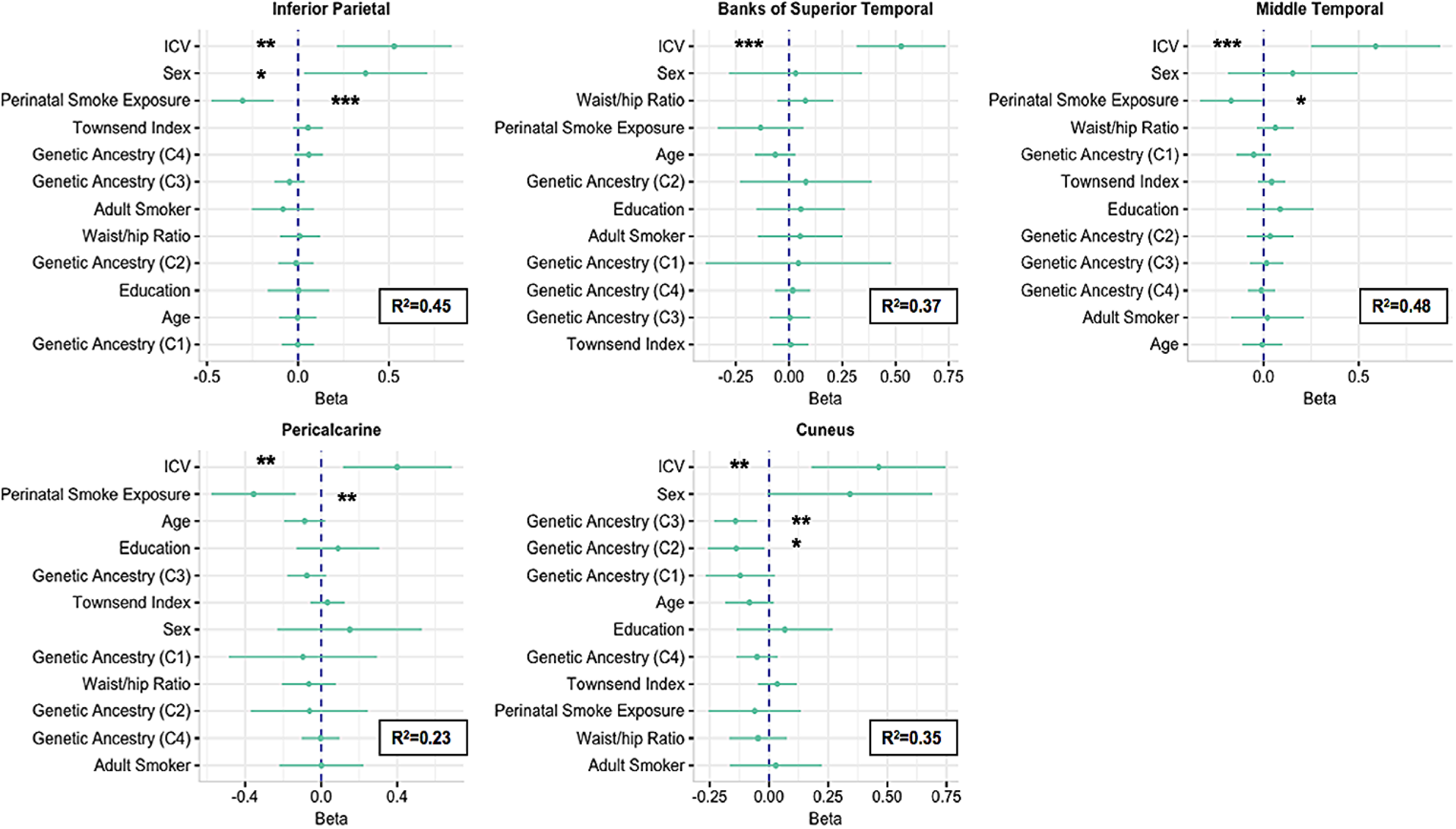
Beta weights of OLS sensitivity analysis in elite healthy replication sample. Predictor variables are plotted on the y axis in order of importance for each model using the lasso method. The individual principle components of genetic ancestry are indicated in parentheses. PSE was not a significant predictor of surface area in the banks of superior temporal sulcus or cuneus. ***p<0.001, **p<0.01, *p<0.05.

Quantile regressions also replicated the findings of lower SA in the PCAL and IPL in the PSE^+^ group, specifically at quantiles 0.5-0.9 and 0.25-0.75, respectively (**Figure 2b, Supplementary Table 3**). We did not replicate the effects of PSE in the bSTS, ITG, MTG, or RMFG, which were observed using the quantile regression method. We did not compute quantile regressions adjusting for birthweight in the elite healthy sample as the design matrix was computationally singular after adjusting for additional covariates.

Effect sizes were moderate in comparison to the discovery sample, particularly in the IPL (unequal variances: PSE^-^, Robust *d*=0.35; PSE^+^, Robust *d* =0.38) and PCAL (unequal variances: PSE^-^, Robust *d*=0.4; PSE^+^, Robust *d* =0.38) for OLS models (**Table 2b**). This finding was also demonstrated with the quantile regressions (Quantile shift effects = 0.61), corresponding approximately to Cohen’s *d*’s =0.4 in the IPL and PCAL (**Table 3b**).

**Table 2.** Effect Sizes of Brain Regions that Differed by PSE Using OLS Regression Models

|  |  |  |  |  |  |  |
| --- | --- | --- | --- | --- | --- | --- |
| Region | a. Population Sample |  |  | b. Elite Healthy Sample |  |  |
|  | PSE<sup>-</sup><br>Robust *d* | PSE<sup>+</sup><br>Robust *d* | *EP* | PSE<sup>-</sup><br>Robust *d* | PSE<sup>+</sup><br>Robust *d* | *EP* |
| Inferior Parietal | 0.15 | 0.15 | 0.10 | 0.35 | 0.38 | 0.27 |
| Pericalcarine | 0.17 | 0.18 | 0.12 | 0.40 | 0.38 | 0.27 |
| Banks of Superior Temporal | 0.12 | 0.11 | 0.08 | 0.15 | 0.14 | 0.10 |
| Middle Temporal | 0.07 | 0.07 | 0.05 | 0.25 | 0.17 | 0.18 |
| Cuneus | 0.08 | 0.08 | 0.05 | 0.17 | 0.18 | 0.13 |

*Note.* Robust Cohen's *d* were acquired with the 'akp.effect' function in R. Effect sizes shown here are independent of covariates in the model. Magnitude of effects can be interpreted according to Cohen's guidelines: 0.2=small, 0.5=medium, $\geq 0.5$ =large. Effect sizes of explanatory power (*EP*) were also computed using 'yuen.effect.ci' in R, as this parameter is robust to unequal variances and can be interpreted as a single value. Cohen's *d*=0.2, 0.5, and 0.8 corresponds to *EP*=0.15, 0.35, 0.5, respectively (Wilcox, in press).

**Table 3.** Robust Effect Sizes of Brain Regions that Differed by PSE Using Quantile Regression

|  |  |  |  |  |
| --- | --- | --- | --- | --- |
| Region | a. Population Sample |  | b. Elite Healthy Sample |  |
|  | *Q* effect | Difference score [CI] | *Q* effect | Difference score [CI] |
| Banks of Superior Temporal SA | 0.53*** | 0.11 [0.06, 0.16] | 0.55 | 0.15 [-0.11, 0.40] |
| Caudate VOL | 0.53*** | 0.10 [0.05, 0.15] | 0.55 | 0.22 [-0.10, 0.41] |
| Cuneus SA | 0.52** | 0.08 [0.03, 0.12] | 0.55 | 0.18 [-0.06, 0.42] |
| Inferior Parietal SA | 0.54*** | 0.15 [0.10, 0.20] | 0.61** | 0.36 [0.14, 0.59] |
| Inferior Temporal SA | 0.52* | 0.05 [0.00, 0.10] | 0.54 | 0.16 [-0.06, 0.38] |
| Middle Temporal SA | 0.52* | 0.07 [0.15, 0.12] | 0.57 | 0.25 [0.01, 0.49] |
| Pallidum VOL | 0.52** | 0.07 [0.02, 0.13] | 0.55 | 0.14 [-0.10, 0.39] |
| Pericalcarine SA | 0.55*** | 0.18 [0.13, 0.23] | 0.61** | 0.14 [0.14, 0.64] |
| Putamen VOL | 0.49 | -0.04 [-0.09, 0.01] | 0.54 | 0.18 [-0.08, 0.43] |
| Rostral Middle Frontal SA | 0.52* | 0.06 [0.01, 0.12] | 0.55 | 0.16 [-0.09, 0.41] |

*Note.* Effect sizes were derived using Yuen's test for trimmed means ('yuenQS' function in R). A robust measure of 'shift', or quantile effect (*Q*) was computed and can be interpreted as follows: Cohen's $d=0, 0.2, 0.5$ , and $0.8$ corresponds to $Q=0.5, 0.55, 0.65$ , and $0.70$ , respectively (Wilcox, in press). Significant *Q* values indicate that we can reject the null hypothesis of equal distribution. *Q* values indicate the quantile to which the distribution has shifted from the population median (0.5). \*\*\* $p<0.001$ , \*\* $p<0.01$ , \* $p<0.05$

### Disease risk by PSE and brain structure

Logistic regressions revealed significant variability in disease risk for each predictor variable in our partial regression models. PSE was associated with increased risk of developing a CNS or psychiatric condition (RR=29%), disease of the ear or mastoid process (RR=63%), circulatory condition (RR=12%), digestive condition (RR=8%), or genitourinary condition (RR=14%) (**Supplementary Table 4**). To determine whether the degree of neuroanatomical insult further contributed to disease risk, we examined the relative risk in the PSE^+^ group after individually adding brain metrics that differed by PSE (SA in the PCAL, IPL, MTG, bSTS, ITG, RMFG, CUN) to the logistic models. Partial *F* tests revealed significantly better fit for each ICD-10 model with the addition of at least one SA metric to the predictor set (**Supplementary Table 5**). However, SA metrics themselves were not always significant predictors of the target condition when also adjusting for all other predictors. For brevity, we describe the models in which regional SA metrics significantly improved model fit *and* significantly predicted the target condition at the 0.05 alpha level.

Analyses in the PSE^+^ group revealed an 11% decreased risk for circulatory conditions (*p*=0.008), and a 10% decreased risk for digestive (*p*=0.002) and musculoskeletal conditions (*p*=0.038) with each standard deviation increase in PCAL SA (**Supplementary Table 6**). We did not observe these relationships in the PSE^-^ group for circulatory or digestive conditions. Surprisingly, we did observe a 7% higher risk for musculoskeletal conditions with each standard deviation increase in PCAL SA in the PSE^-^ group (*p*=0.042). The PSE^+^ group also demonstrated an 11% decreased risk for circulatory and genitourinary conditions with each standard deviation increase in SA of the MTG and RMFG, respectively. Finally, SA in the IPL, ITG, and MTG was associated with a 19%, 16%, and 15% decreased risk for endocrine conditions in the PSE^+^ group with each standard deviation increase in SA. These risks were not observed in the PSE^-^ group. Across all logistic regressions where brain metrics improved model performance, the addition of the IPL as a predictor of endocrine conditions in the PSE^+^ group showed the best model fit as determined by Akaike information criterion (AIC).

For interpretation, we examined the role of subclasses of the ICD-10 conditions identified above in relation to PSE. Greater SA in the PCAL and MTG was associated with lower risk for hypertensive disease in the PSE^+^ (RR=0.87 and 0.82, respectively) and PSE^-^ groups (RR=0.90 and 0.83, respectively). Lower SA in the PCAL also was associated with a higher risk for non-infective gastroenteritis/colitis (RR=0.72), colon polyps (RR=0.63), and joint disorders (RR=0.81) in the PSE^+^ group only. Finally, lower SA in the RMFG was associated with higher risk for pelvic inflammatory disease (RR=0.56), and endometrial polyps (RR=0.50) among females in the PSE^+^ group only (**Supplementary Table 7**). We did not observe a significant effect of SA on sub-classes of endocrine conditions in either PSE group.

## Discussion

We report the first large-scale evidence of long-term cortical brain disruption among individuals perinatally exposed to tobacco smoke. Consistent with our hypothesis, cortical SA measurements were more sensitive to PSE than CT or VOL. PSE^+^ individuals demonstrated lower SA in occipito-parietal and temporal brain regions when compared to the PSE^-^ group, with the most robust effects observed in the pericalcarine (PCAL) and inferior parietal cortex (IPL). Cortical SA abnormalities in the PCAL, middle temporal gyrus (MTG), and rostral middle frontal gyrus (RMFG) were associated with significantly higher risk for disease among individuals in PSE^+^ group, with the PCAL affecting risk for a broad set of disorders within the circulatory, digestive, and musculoskeletal systems. Importantly, lower SA in the PCAL and IPL was observed after adjusting for potential psychosocial and physiologic confounders in both the population-based sample and a hold-out replication sample of healthy adults, suggesting the observed relationships to PSE were not a downstream effect of intervening health conditions. Collectively, these findings highlight the long term consequences of PSE on brain structure and disease risk later in life.

Cortical SA was most closely associated with PSE - consistent with the developmental timing of brain structure maturation in the perinatal period. Cortical maturation is sensitive to environmental exposures during this time, as the majority of cortical growth occurs between 21 weeks’ gestational age and the first postnatal year (Gilmore et al. 2012). However, SA and CT follow distinct patterns of pre- and postnatal development and show consistently unique relationships with age. The majority of CT growth occurs during gestation, followed by an increase in CT around 40% and 0.7% during the first and second postnatal years, respectively. Changes in CT during the first postnatal year have been shown in nearly all regions except the central sulcus and calcarine fissure, with the lowest growth rates in visual, motor, and somatosensory cortices (Li et al. 2015). By contrast, the cortical surface expands by around 180% and 120% during the first and second postnatal year, with localized gains in temporal, parietal, and occipital cortices (Li et al. 2013). More recent work shows a 0.51% daily increase in SA and 0.09% daily increase in CT between 6 and 144 days after birth (Jha et al. 2018). Changes in cortical development after age 2 are primarily driven by cortical surface expansion (Lyall et al. 2015), suggesting there is an extended window of vulnerability of cortical SA to secondhand smoke and other environmental toxins beyond the perinatal period. This is consistent with several reports of monotonic decline of CT after age 3, though this pattern has not been universally identified across studies (for review see Walhovd et al. 2017). Data related to childhood and adolescent smoke exposure were not available for our study, but it is possible that our results reflect chronic effects of secondhand smoke that occurred throughout childhood development. Future studies will benefit from a more detailed assessment of early life environment that includes information on duration of exposure, frequency of smoking behaviors and number of smokers in the household.

The link between PSE and lower SA in the PCAL and IPL is consistent with animal studies that revealed altered brain development in the visual and somatosensory systems following prenatal nicotine exposure. The most prominent nAChR subunit in the human CNS is heteromeric α4β2, which binds nicotine with high affinity (Bansal et al. 2000). Earlier work using autoradiography in rats revealed a 90% and 107% increase in α4β2 density in the somatosensory and visual cortices, respectively, following prenatal nicotine administration (Tizabi and Perry 2000). Primate studies by Duncan et al. (2009; 2015) reported increased nAChR binding, including α4β2, in the medulla and primary visual cortex (V1) of fetal baboons. Receptor upregulation in these regions is believed to reflect the process of desensitization, increasing the risk for developmental perturbations (Ernst et al. 2001; Aoyama et al. 2016). Early postnatal smoke exposure also alters normal programming of the cholinergic system (Navarro et al. 1989; Yanai et al. 1992; Zahalka et al. 1992; Slotkin 2004; Heath and Picciotto 2009), which prior studies have linked to altered developmental plasticity, reduced cortical size, and attenuated maturation of cortical pyramidal neurons (Aramakis and Metherate 1998; Robertson et al. 1998). Developmental disruptions in cholinergic activity may explain lower cortical SA in the PCAL of exposed individuals in our study, as spectroscopy has revealed elevated choline in the PCAL of subjects with congenital blindness - a contrast to typically high choline levels at birth that decline with postnatal cortical maturation and pruning (Bluml et al. 2013). We suspect that abnormal developmental programming associated with PSE may be driven by epigenetic modifications that alter the biological response to stress and arrest normal brain maturation (Knopik et al. 2012). Epigenetic changes were not investigated here, but maternal tobacco use during pregnancy may disrupt epigenetic regulation by changes in DNA methylation (Toledo-Rodriguez et al. 2010; Ba et al. 2011; Suter et al. 2011). Future studies examining longitudinal DNA methylation patterns among individuals exposed to PSE will clarify this possible connection.

Negative associations between SA in the PCAL and MTG and risk for circulatory conditions in the PSE^+^ group extends prior work. In Tchistiakova et al. (2014), individuals with hypertension and type 2 diabetes demonstrated lower cerebrovascular reactivity in occipitoparietal cortices during successive breath holds, and lower thickness in the right occipital lobe. Autopsy analysis also has shown microinfarcts in the occipital cortex of brains with cerebral amyloid angiopathy – a cerebrovascular condition characterized by amyloid build-up in blood vessels of the CNS (Kovari et al. 2017). Prior work using structural MRI has revealed positive associations between CT in the MTG and cholesterol (Leritz et al. 2011), further supporting a role for vascular health on this brain region.

The link between SA in the PCAL and musculoskeletal conditions was not expected, but may be partly explained by increased proteolysis through smoke-induced activation of inflammatory mediators. Exposure to cigarette smoke increases blood levels of proinflammatory cytokines such as TNF-α and IL-6 in animals, which can lead to muscle wasting by increased proteolysis and protein synthesis inhibition (Degens et al. 2015). Acetaldehyde, a Group 2B substance (*possibly carcinogenic*) and bioactive toxicant of mainstream and sidestream smoke, lowers the rate of protein synthesis in cultured human muscle cells (Hong-Brown et al. 2001). Excessive proteolysis via acetylaldehyde activation of matrix metalloproteins (MMPs) is sufficient to destroy articular cartilage in diseased joints (Rengel et al. 2007; Rose and Kooyman 2016) and may explain the increased risk for joint disorders in the PSE^+^ group. Imbalanced proteolytic activity is also implicated in digestive conditions, as MMPs are the most abundant proteases in gut mucosa and are upregulated among individuals with ulcerative colitis (Pujada et al. 2017). While these mechanisms offer potential insights into links between PSE and disease, increased risk for these disorders as a function of the PCAL is not clear. Smoke-induced exposure to heavy metals (Otto and Fox 1993; Korogi et al. 1994; Braissant et al. 2013; Ekinci et al. 2014; Oeltzschner et al. 2015) is associated with biochemical disruptions to neurotransmitter systems, and it is possible that physiologic and neurostructural changes explain the link between the visual cortex and peripheral diseases of the digestive and musculoskeletal systems. Alternatively, the degree of SA disruption in the PCAL may be a marker of extant exposure, and there may not be a direct link between PSE and PCAL structure directly. Experimental studies are needed to determine the degree that toxins from secondhand smoke beyond nicotine influence central and peripheral biochemistry.

This study has several strengths. First, the large sample size boosted statistical power and allowed us to create a hold-out sample of healthy adults to test the reproducibility of our main findings. Replication analyses revealed a specific effect of PSE on the PCAL and IPL; stronger effect sizes in these regions in the healthy hold-out sample may indicate a more direct effect of PSE than other brain regions identified in the main analyses. Importantly, the replication sample was on average 3 years younger than the population-based sample and may be at high risk for PCAL-associated diseases of the circulatory and digestive system with increasing age (PCAL-associated risk for musculoskeletal conditions did not show an aging effect). Although the IPL was not identified as a significant predictor of disease risk in this sample, SA in this region did improve model fit for diseases of the circulatory and respiratory systems. The lack of direct link may be due to low cell sizes for specific conditions within these larger ICD-10 categories.

Second, we used robust statistical methods to ensure the most accurate depiction of PSE brain effects. Part of this approach included the use of quantile regression models to examine the effect of PSE at various points along the distribution. By doing so, we observed additional region-specific effects of PSE in the ITG and RMFG - regions that were not captured using a standard regression approach. SA in these regions improved model fit for our logistic regression models of CNS and genitourinary conditions, respectively. The ITG did not show a significant individual relationship to CNS conditions, but higher SA in the RMFG was linked to a 44-50% lower risk of pelvic inflammatory disease (PID) and endometrial polyps in females of the PSE^+^ group. This association may be partly explained by the increased risk for PID with increasing cardiovascular morbidities (Chen et al. 2011), the latter showing links to cerebral hypoperfusion in the RMFG (Hoscheidt et al. 2017). Of note, recent histological analysis of the human uterine cavity revealed positive associations between endometrial polyps and heavy metals such as lead, cadmium, aluminum, and nickel – trace elements found in tobacco smoke (Rzymski et al. 2016).

A few limitations of this study should be noted. First, our measure of PSE was nonspecific, and the developmental timing and duration of exposure is unknown. Information on paternal smoking around birth and during childhood was not available, nor was information on additional parental health behaviors that could affect brain development, particularly maternal alcohol consumption. Alcohol contains many of the same toxins as cigarette smoke (e.g., aldehydes) and maternal alcohol consumption during pregnancy imparts a similarly high risk to fetal brain development (Kitsiou-Tzeli and Tzetis 2017). Still, it is unlikely that light to moderate prenatal alcohol exposure was a primary driver of our results since the window of exposure is limited (9 months) when compared to the potential window of tobacco exposure that imposes high risk to brain development (through adolescence and beyond). Further, we attempted to account for these limitations by conducting a secondary analysis that covaried for birth weight, which prior studies have identified as a stable marker of prenatal health (Gortmaker 1979; Kogan et al. 1994; Alexander and Korenbrot 1995). These analyses revealed a nominal change in results, suggesting that our results were not confounded by other prenatal health factors. Second, smoking initiation and regular smoking behaviors are heritable traits (for review see Do and Maes 2017), so participants whose mothers smoked may have inherited a greater aggregate risk from genetic variants that predispose to smoking initiation and persistence among smokers in this study compared to those whose mothers did not smoke. As such, the effect of PSE on imaging derived brain measures may partly result from smoking-associated genetic variants that were inherited among exposed participants. Third, we cannot assess the directionality of disease risk for each brain measure in our logistic models given the descriptive nature of this design. Finally, we do not yet know if our results generalize to other populations, as this sample was specific to a UK population and disease prevalence in the UKB sample is lower, on average, than disease prevalence in other population-based cohorts (Fry et al. 2017).

## Conclusion

This is the largest-ever neuroimaging study of the structural brain effects of PSE, and the first study in a sample of middle and older-aged individuals. Our imaging and robust statistical approach revealed a dominant, system-specific effect of PSE on sensory brain structures later in life, with the most pronounced and clinically relevant abnormalities on cortical SA of the PCAL. The findings in this study are of high public health relevance given the current international tobacco crisis that is increasing in some low and middle income countries (WHO 2018). Given global increases in life expectancy, long-term effects of PSE may bring added morbidities and complicate quality of life in later adulthood. Further, the evolving landscape of cannabis legislation may pose many of the neuroanatomical consequences and overall health risks discussed here as a result of smoke exposure (Moir et al. 2008).

## Acknowledgments

This work was conducted using the UK Biobank Resource under Application Number 11559. The UK Biobank was established by the Wellcome Trust, Medical Research Council, Department of Health and the Scottish Government. UK Biobank funding was also supported by the Welsh Assembly Government, British Heart Foundation, and Diabetes UK. Cortical processing and QC were supported by the ENIGMA pipeline (http://enigma.ini.usc.edu/protocols/imaging-protocols/).

## Footnotes

^1^A levels= Advanced level subject-based qualification as part of the General Certificate of Secondary Education (GCSE); O levels/GCSEs = Ordinary Level GCSE; CSE= Certificate of Secondary Education; NVQ= National Vocational Qualification; HND= Higher National Diploma; HNC= Higher National Certificate “

